# The experimental design and data interpretation in “Unexpected mutations after CRISPR–Cas9 editing *in vivo*” by Schaefer et al. are insufficient to support the conclusions drawn by the authors

**DOI:** 10.1101/153338

**Authors:** Christopher J. Wilson, Tim Fennell, Anne Bothmer, Morgan L. Maeder, Deepak Reyon, Cecilia Cotta-Ramusino, Cecilia A. Fernandez, Eugenio Marco, Luis A. Barrera, Hariharan Jayaram, Charles F. Albright, Gerald F. Cox, George M. Church, Vic E. Myer

## To the Editor

The recent correspondence to the Editor of Nature Methods by Schaefer et al.^1^ has garnered significant attention since its publication as a result of its strong conclusions contradicting numerous publications in the field using similar analytical approaches and methods^2–4^. The authors suggest that the CRISPR-Cas9 system is highly mutagenic in genomic regions not expected to be targeted by the gRNA. We believe that the conclusions drawn from this study are unsubstantiated by the disclosed experiments as they were designed and carried out. Further, it is impossible to ascribe the observed differences in the subject mice to the effects of CRISPR *per se*. The genetic differences seen in this comparative analysis were likely present prior to editing with CRISPR.

In our view, the experiments, observations, and subsequent assertions in Schaefer et al.^1^ can be summarized as follows. Two mice created using CRISPR-based genome editing at the zygote stage, when compared to a single “co-housed FVB/NJ mouse without CRISPR-mediated correction”, showed a significant number of single nucleotide variants (SNVs) and insertions and deletions (indels) across the genome. The number of mutations common to the two independently generated CRISPR edited mice was 1,397 SNVs and 117 indels. Surprisingly, these apparent mutations all arose at regions in the genome that have poor homology to the gRNA (between 5% – 65%). Furthermore, none of the 50 closest, predicted off-target sites (based on gRNA sequence homology) had any observed activity (SNVs or indels). The authors speculate that there is an unreported activity where “certain sgRNAs may target loci independently of their target *in vivo*.”

Our opinion is that the conclusions drawn from this study are unsubstantiated by the disclosed experiments and that it is impossible to ascribe the observed differences in the subject mice to the effects of CRISPR *per se* based upon the following observations:

Firstly, the overall number of the study subjects is low (*n* = 2 treated mice and *n* = 1 untreated mouse) and the sequencing depth applied to the treated and untreated mice is not equivalent. An underpowered study may prove limiting when attempting to understand statistical reproducibility and reliability of scientific observations.

Secondly, the selection of a co-housed mouse (as opposed to the parents or *bona fide* littermates) as the control is insufficient to attribute the observed differences between the treated mice and control mouse to CRISPR. The design of the experiment makes it impossible for the authors to rule out the possibility that the reported genomic differences between the experimental animals and the single control existed prior to experimental manipulation with CRISPR. In fact, published literature has shown that differences in the genomes of *littermates* analyzed by whole genome sequencing (WGS) can be significant (985 SNVs were identified by Oey et al.^5^). These differences are attributed to private mutations propagated by normal Mendelian inheritance within a breeding colony. In Oey et al., further analysis of the parents by sequencing methods confirmed the vast majority of these SNVs were present in the parents and a small minority arose as private variations in the progeny^5^.

To further understand the observations in Schaefer et al.^1^ we reanalyzed their sequencing data deposited in the NCBI-SRA database. Raw sequence (fastq) files were retrieved, and, because the analysis parameters were not sufficiently described to reproduce the authors’ analysis, we re-aligned and identified variants using a standard analytical framework described in the supplement to this letter. Similar to Schaefer et al. we identified SNV and indel differences between the control “FVB” mouse and the test “F03” and “F05” mice, with 4,022 SNVs and 2,799 indel variants found across the three mice. We focused our initial analysis on variants where there are only two alleles in the three test mice; filtering out variants where there are either three or more alleles across the three mice or all alleles are identical in all mice yet distinct from the mouse reference sequence (mm10); leaving 3,978 SNVs and 2,713 indel variants for analysis (summarized Table 1). Our analysis shows a striking similarity in SNVs that are identical between F03 and F05 but distinct from FVB (2,447). In fact, the frequency of changing both alleles to the same sequence was almost two-fold higher than the frequency of changing either the F03 or F05 mouse alone (874 and 645 alleles respectively). Such a strong similarity between the F03 and F05 mice is unexpected for a random mutagenesis event during the independent creation of these mice, and suggests either underlying genetic similarities or a mutagen that is strongly directive.

**Table 1:**
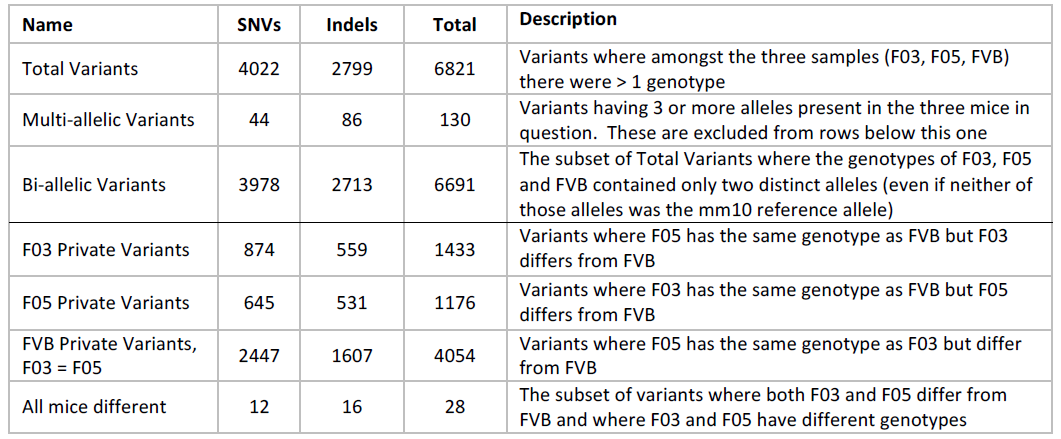
Variant counts

| Name | SNVs | Indels | Total | Description |
| --- | --- | --- | --- | --- |
| Total Variants | 4022 | 2799 | 6821 | Variants where amongst the three samples (F03, F05, FVB) there were > 1 genotype |
| Multi-allelic Variants | 44 | 86 | 130 | Variants having 3 or more alleles present in the three mice in question. These are excluded from rows below this one |
| Bi-allelic Variants | 3978 | 2713 | 6691 | The subset of Total Variants where the genotypes of F03, F05 and FVB contained only two distinct alleles (even if neither of those alleles was the mm10 reference allele) |
| F03 Private Variants | 874 | 559 | 1433 | Variants where F05 has the same genotype as FVB but F03 differs from FVB |
| F05 Private Variants | 645 | 531 | 1176 | Variants where F03 has the same genotype as FVB but F05 differs from FVB |
| FVB Private Variants, F03 = F05 | 2447 | 1607 | 4054 | Variants where F05 has the same genotype as F03 but differ from FVB |
| All mice different | 12 | 16 | 28 | The subset of variants where both F03 and F05 differ from FVB and where F03 and F05 have different genotypes |

When reviewing the variant list, we included annotation as to whether the variant was found in the mouse reference genome (mm10), a Black 6 strain. It immediately became obvious that many of the variants are distributed relative to the mm10 reference in a way that would not be expected if a mutagen were applied (like CRISPR/Cas9, as proposed by the authors, or potentially another step in the process). For example, as summarized in Table 2, there are 2,508 SNVs where the FVB mouse genotype is homozygous and matches the mm10 reference and the F03 or F05 mice have a different genotype. Of these, 409 (16%) are “complete switches”, where the F03 and F05 have identical homozygous genotypes that are not the mm10 reference. However, when examining the 730 SNVs in the FVB control mouse that are homozygous for a genotype not matching the mm10 reference, a striking 578 SNVs (79%) appear as “complete switches” for both the F03 and F05 mice *back* to the homozygous mm10 reference. Additionally, there are only 27 variants (4%) where both F03 and F05 mice have homozygous changes that do not match the mm10 reference. When considering just “complete switches,” an expected distribution of SNVs would be 66% to one of the two non-mm10 references and 33% to the mm10 reference, yet here we see 4% and 96% respectively – making this deviation highly significant (Chi-Squared p<0.00001). An analysis with indels yields similar results. Of 1,698 homozygous indels matching mm10 in the FVB mouse 458 are “complete switches” (27%) in F03 and F05, and of 779 homozygous non-mm10 indels in FVB, 285 (36%) are complete switches back to the mm10 reference. However, only 126 (16%) are complete switches to another genotype. It is impossible to calculate an expected distribution because the number of possible indels is much larger and not defined. However, there is no reason to expect that indels would appear with a greater than two-fold preference for the reference mm10 sequence over any other possible indel. The SNV and indel analyses for these extreme “full switch” scenarios indicate that a mutagen (either CRISPR/Cas9 or other process steps) is unlikely to be causative for these observed variants, and, with such a strong signal relative to the mm10 reference, it argues for an alternate explanation including variation in the breeding colony and subsequent Mendelian inheritance.

**Table 2:** Analysis of variant counts

|  | SNVs | Indels |
| --- | --- | --- |
| FVB mouse homozygous and genotype **matches mm10** | 2,508 | 1698 |
| "Complete switches" where F03 and F05 have the same genotype and are different than FVB (**not matching mm10**) | 409 (16%) | 458 (27%) |
| All other F03 and F05 genotypes in this | 2,099 (84%) | 1,240 (73%) |
| FVB mouse homozygous and genotype **does not match mm10** | 730 | 779 |
| "Complete switches" where F03 and F05 have the same genotype **matching mm10** | 578 (79%) | 285 (36%) |
| "Complete switches" where F03 and F05 have the same genotype **not matching mm10** | 27 (4%) | 126 (16%) |
| All other F03 and F05 genotypes in this set | 125 (17%) | 368 (47%) |

Heterozygosity was called out in a recent letter posted on BioRxiv (http://dx.doi.org/10.1101/154450) from the Schaefer et al. authors as unexpected and differential between the F03 & F05 and the FVB. To better understand these claims, we went on to examine heterozygosity within the three mice described in Schaefer et al^1^. and, for comparison, we included the FVB/NJ mouse sequenced at an average of 50x coverage by the Sanger Center as part of the Mouse Genome Project^6^. Strikingly, we find high levels of heterozygosity (˜144,000 SNV het calls per mouse) at roughly equal levels in all four mice. This high heterozygosity is dependent on turning off filters that remove variants with exceptionally high read coverage. Applying these filters reduces the heterozygous counts a little less than 10-fold (˜17,000 SNV het calls per mouse), but all four mice still have roughly equivalent levels of heterozygosity (Table 3). While there are clearly some heterozygous calls that differ between the mice (as pointed out by Schaefer et al.), as shown in Figure 1, most of the heterozygous SNVs (>85%) are shared between all four mice. The Schaefer et al. statements that there is excess heterozygosity in the F03 and F05 is simply not supported in our reanalysis of the data.

**Table 3:** Analysis of heterozygosity

| Type | Mouse | Including High Coverage Reads | Excluding High Coverage Reads |
| --- | --- | --- | --- |
| SNVs | F03 | 144,633 | 17,097 |
|  | F05 | 143,766 | 16,827 |
|  | FVB | 143,413 | 15,864 |
|  | FVB/NJ (Sanger) | 145,439 | 17,723 |
| Indels | F03 | 15,710 | 2,650 |
|  | F05 | 15,684 | 2,669 |
|  | FVB | 15,108 | 2,013 |
|  | FVB/NJ (Sanger) | 15,724 | 2,554 |

**Figure 1:**
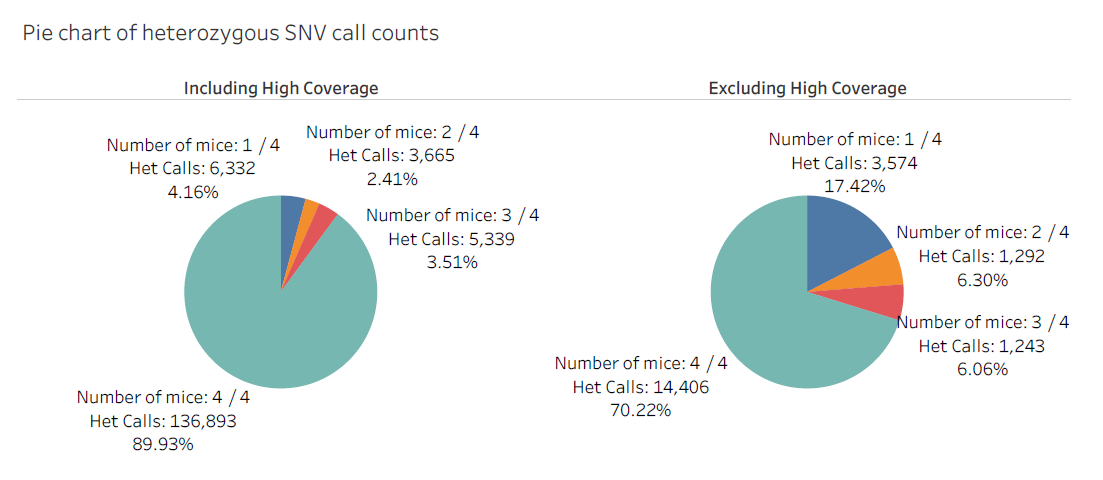
Pie charts illustrating the high number of shared SNV heterozygous alleles in the 4 mice either without or with high read count filters applied.

The finding that there are a large number of heterozygous SNVs stably inherited within the inbred FVB/NJ mouse strain is interesting in its own right. One hypothesis is that these SNVs are found in nearly identical duplicated regions of the genome. Thus, what appears to be a heterozygote at one locus, may, in fact, be two almost identical duplicated genomic regions that differ only by that one nucleotide. The fact that high coverage filters reduces these heterozygous calls by ˜90% supports this notion of duplicated regions. It is also consistent with the inbreeding procedures used to maintain the FVB/NJ mouse colonies, where heterozygosity at this scale is unexpected, as Schaefer et al. point out. Other explanations for the high level of heterozygosity may also emerge with more analysis or sequencing. These findings highlight the notion that genomes are more complex and not as well characterized, especially for the mouse, than often perceived.

Improvements in experimental design would greatly improve the Schaefer et al. study. In order to control for the reality that inbred mice are not perfectly identical at the nucleotide level and our understanding of their genomes in incomplete, an appropriately controlled experiment would include essential components such as 1) sequencing of the parent animals to ascertain the input genome sequences going into the experiment, 2) breeding out the CRISPR edited mice to remove chimerism, and 3) generating and characterizing mice using identical methodology derived from the same experimental protocol, but lacking key individual components, to rule out the possibility that the method itself was mutagenic. More specifically, mice generated with plasmid (encoding the sgRNA) + single stranded DNA oligonucleotide (ssODN) donor DNA +Cas9 protein should be compared to mice generated with plasmid + ssODN donor, plasmid + Cas9 protein, and ssODN donor + Cas9 protein. This would control for the possibility that either of these components individually, or the process of generating the mice, was inherently mutagenic. A similar study^2^ has been published in the same journal using appropriate controls and finding significantly lower SNVs and indels suggesting experimental differences, and not CRISPR, are likely causes of the recent observations of Schaefer et al. ^1^

Furthermore, we would highlight the following observations reported in the Schaefer et al. ^1^ communication:

The specific gRNA used in the disclosed experiments, when run through gRNA specificity prediction algorithms, shows a high propensity for off targets, identifying 1 off-target site that differs from the mouse genome by 1 nucleotide match, 1 off-target site that differs from the mouse genome by 2 nucleotide matches, and 24 off-target sites that differ from the mouse genome by 3 nucleotide matches. While perhaps acceptable for research purposes, a gRNA with a predicted high off-target profile would be immediately excluded as a therapeutic candidate. Despite the high propensity for off target activity we found it surprising that this gRNA showed none of the predicted off-targets using the methods employed in this study underscoring the importance of both predicting and testing empirically for off-target activity.

To underscore potential phenotypic consequences, Schaefer and coauthors focused on an analysis of exonic changes. Most exonic SNVs found in the two CRISPR edited mice (Schaefer et al.^1^, Supplemental Tables 1 and 2) were not only shared between these mice, despite the assertion that the SNVs were created in separate zygotes, but also exhibited identical nucleotide changes in both position and nucleotide composition. Both animals were either homozygous or heterozygous for the same nucleotide change at the same genomic position. As highlighted by our analysis as well, *this strongly suggests the vast majority of these mutations were present in the animals of origin. The odds of the exact nucleotide changes occurring in the exact same position of the exact same gene in almost every case are effectively zero.*

To summarize, our opinion is that the authors failed to sufficiently control the reported study in such a way that one could conclude that CRISPR induces the observed mutations. In our view, the genetic differences seen in this comparative analysis were likely present prior to editing with CRISPR. We encourage the authors to follow up with an appropriately controlled experiment as understanding and controlling the specificity of CRISPR technology is essential for research and critical for therapeutic development. We are firmly committed to a rigorous, objective, and comprehensive assessment of specificity in our own work and seek to advance a shared understanding in the field of how to best assess this critical parameter for bringing CRISPR-based medicines to patients with genetically-defined or genetically-treatable diseases.

## Conflict of Interest Statement

AB, MLM, DR, CJW, CCR, CAF, EM, LAB, HJ, CFA, GFC and VEM are full time employees of Editas Medicine. TF is a paid consultant of Editas Medicine. GMC is an advisor to Editas Medicine and his competing interests are covered here: http://arep.med.harvard.edu/gmc/tech.html

## Supplemental Information

Sequence data for F05 (SRR5450996), F03 (SRR5450997) and FVB (SRR5450998) was retrieved from the Short Read Archive and converted to FASTQ format. Data was processed through a pipeline consisting of a) realignment to the GRCm38/mm10 reference genome using bwa-mem^s1^ (version 0.7.15-r1140), b) duplicate removal (FVB/NJ PCR+ sample only) using Picard’s MarkDuplicates (version 2.9.2), c) variant detection and joint-genotyping using the GATK^s2^ HaplotypeCaller (version 3.7-0-gcfedb67) and d) variant filtration. Full command lines are given in Supplementary Table 1.

**Supplementary Table 1.**

| Task | Command Line |
| --- | --- |
| Low Complexity Masking | minimap/sdust mm10.fa > low_complexity.bed |
| Alignment | bwa mem -t \$threads -p mm10.fa sample.fq |
| Duplicate Marking | java picard.jar MarkDuplicates CREATE_INDEX=true I=\$sample.bam O=\$sample.deduped.bam M=\$sample.metrics.txt |
| Variant Discovery | java -Xmx4096m -jar GenomeAnalysisTK.jar -T HaplotypeCaller -R mm10.fa -L regions.bed --minPruning 3 --maxNumHaplotypesInPopulation 200 --emitRefConfidence GVCF --max_alternate_alleles 3 --contamination_fraction_to_filter 0.0 -I \$sample.bam -o \$sample.g.vcf.gz -pairHMM VECTOR_LOGLESS_CACHING |
| Joint Genotyping | java -Xmx4096m -jar GenomeAnalysisTK.jar -T GenotypeGVCFs -R mm10.fa -L regions.bed --dbsnp dbsnp146.mm10.vcf.gz -V F03.g.vcf.gz -V F05.g.vcf.gz -V FVB.g.vcf.gz -V FVB_NJ.g.vcf.gz -o calls.vcf.gz |
| Filtration | java -Xmx4g -jar picard.jar FilterVcf MIN_AB=0.3 I=calls.vcf.gz O=filtered.vcf.gz CREATE_INDEX=true |

Variant calling was restricted to autosomal regions that were not identified as low-complexity by sdust (approximately 6% of autosomal sequence is identified as low complexity).

Variant calls were filtered to provide a high quality set of variant calls for analysis. Picard’s FilterVcf was used to filter out variants with heterozygous genotypes where either allele accounted for < 30% of the observations in heterozygous samples. Custom filters were applied to remove variants where a) any sample was unable to be genotyped or had less than 23X coverage of the variant, b) any sample had exceptionally high coverage (defined as coverage greater than the sample mean plus three times the square root of the sample mean), c) all samples shared the same genotype, or d) more than two alleles were observed across all samples.

## References

1. Schaefer, K. A. et al.. Unexpected mutations after CRISPR–Cas9 editing *in vivo*. 1–2 (2017). doi:10.1038/nmeth.4293

2. Iyer, V. et al.. Off-target mutations are rare in Cas9-modified mice. Nature Methods 12, 479–479 (2015).

3. Smith, C. et al.. Whole-Genome Sequencing Analysis Reveals High Specificity of CRISPR/Cas9 and TALEN-Based Genome Editing in Human iPSCs. Cell Stem Cell 15, 12–13 (2014).

4. Veres, A. et al.. Low Incidence of Off-Target Mutations in Individual CRISPR-Cas9 and TALEN Targeted Human Stem Cell Clones Detected by Whole-Genome Sequencing. Cell Stem Cell 15, 27–30 (2014).

5. Oey, H., Isbel, L., Hickey, P., Ebaid, B. & Whitelaw, E. Genetic and epigenetic variation among inbred mouse littermates: identification of inter-individual differentially methylated regions. Epigenetics & Chromatin 1–12 (2015). doi:10.1186/s13072-015-0047-z

6. Wong, K et al.. Sequencing and characterization of the FVB/NJ mouse genome. Genome Biology 13:R72 (2012).

## References

S1. Li, Heng. Aligning sequence reads, clone sequences and assembly contigs with BWA-MEM. arXiv:1303.3997.

S2. McKenna, A. et al.. The Genome Analysis Toolkit: a MapReduce framework for analyzing next-generation DNA sequencing data. doi:10.1101/gr.107524.110.

